# Inhibition of *Pf*CLK3 a master regulator of malaria parasite RNA-splicing provides the basis for a radical cure for malaria

**DOI:** 10.1101/2025.10.07.680601

**Authors:** Omar Janha, Niniola Olaniyan, Go Ka Diam, Saumya Sharma, Dario Beraldi, Andrew G. Jamieson, Zbynek Bozdech, Andrew B. Tobin, Katarzyna Modrzynska

## Abstract

Emerging resistance to front-line anti-malarials means there is a race to discover new drugs with novel mechanisms of action that remain effective across multiple stages of the parasite life cycle. Previously we reported the malaria protein kinase, *Pf*CLK3, as a target offering a cure, prophylaxis and transmission blocking. The homology between *Pf*CLK3 and human kinases suggested that the mechanism of parasiticidal activity of *Pf*CLK3 inhibitors is disruption of RNA processing. Here, we use whole genome RNA-sequencing to reveal that selective *Pf*CLK3 inhibition extensively affects RNA-splicing with 2039 splice-junctions across 1125 genes mis-spliced in treated wild type parasites compared to controls. The function of mis-spliced transcripts showed that the affected genes were involved in numerous essential parasite processes associated with multiple life cycle stages and revealed transcripts and introns particularly susceptible to inhibition. Our study supports the role of *Pf*CLK3 as an important regulator of spliceosome activity and establishes the distinct mechanism of parasiticidal activity of *Pf*CLK3 inhibitors from current front-line treatments.

## Introduction

Despite significant gains in the fight against malaria, progress in the last two decades has stalled, partly due to the emergence and spread of parasites resistant to the front-line antimalarial drug, artemisinin, and its partners in artemisinin combination therapies (ACTs). Hence, new antimalarial drugs with alternative modes of action against novel targets are critically needed (1). Ideally, these novel targets are conserved across parasite species, especially those bearing the highest impact, *P. falciparum* and *P. vivax* (2).

It is therefore of interest that among the 36 eukaryotic protein kinases that have been reported to be essential for the survival of the asexual blood stage of the most virulent species of human malaria, *P. falciparum*, the cyclin-dependent like protein kinase family (CLK family) made up of 4 members (*Pf*CLK1-4) (1,2) have emerged as potential drug targets. Prominent among this kinase family is *Pf*CLK3 which is highly conserved amongst *Plasmodium spp* and its inhibition has been shown to have multi-stage and cross species parasiticidal activity (1). In this way we and others have suggested that development of drugs inhibiting *Pf*CLK3 would have the potential to offer a cure, prophylaxis and transmission blocking medicine that is active across all human *Plasmodium spp*.

The similarity of the *Pf*CLK-family to the mammalian CLKs and the serine-arginine-rich protein kinase (SRPK) family implies the ability of *Pf*CLK3 to phosphorylate SR proteins (2–5). Additionally, the kinase catalytic domain of *Pf*CLK3 is closely related to the human splicing factor kinase PRP4 kinase (PRPF4B) (6–8) further supporting a role for this kinase in RNA-processing (5). Initial studies from our laboratory have reported that selective inhibition of *Pf*CLK3 down-regulates RNA transcribed from genes that are essential for parasite survival (1). Despite these early studies and an advanced medicinal chemistry programme developing inhibitors to *Pf*CLK3 there has been no direct investigation of *Pf*CLK3’s role in RNA processing.

Here we present total RNA-sequencing data following *Pf*CLK3 inhibition in wild type (WT), and genetically engineered parasites expressing an inhibitor resistant variant of *Pf*CLK3, G449P, in a study designed to investigate the impact on RNA processing (9). Our studies show that specific *Pf*CLK3 inhibition results in extensive disruption of RNA splicing in genes encoding proteins involved in essential biological processes across multiple stages of the parasite life cycle. In this way, our study confirms *Pf*CLK3 as an important regulator of spliceosome activity and establishes the mechanism of parasiticidal activity of *Pf*CLK3 inhibitors-as disruption of RNA-splicing. It also provides a framework for further investigation of the mechanism of splicing in *Apicomplexa* and optimisation CLK3 selective inhibitors.

## Results

To investigate the role of *Pf*CLK3 in splicing, we first determine conditions under which the *Pf*CLK3 inhibitor TCMDC-135051 could be used to inhibit *Pf*CLK3 activity without causing parasite death, which would otherwise result in extensive secondary changes in transcriptome. We previously determined the EC_50_ of TCMDC-135051 for parasite killing to be approximately 200 nM in a 72-hour drug assay (1). Using this as a benchmark, doubly synchronised trophozoite-stage parasites were exposed to varying concentrations of TCMDC-135051 (0.25 - 4 μM) for 60 minutes to identify an inhibitor concentration that would not adversely affect parasite growth **(Fig. 1A)**.

**Figure 1:**
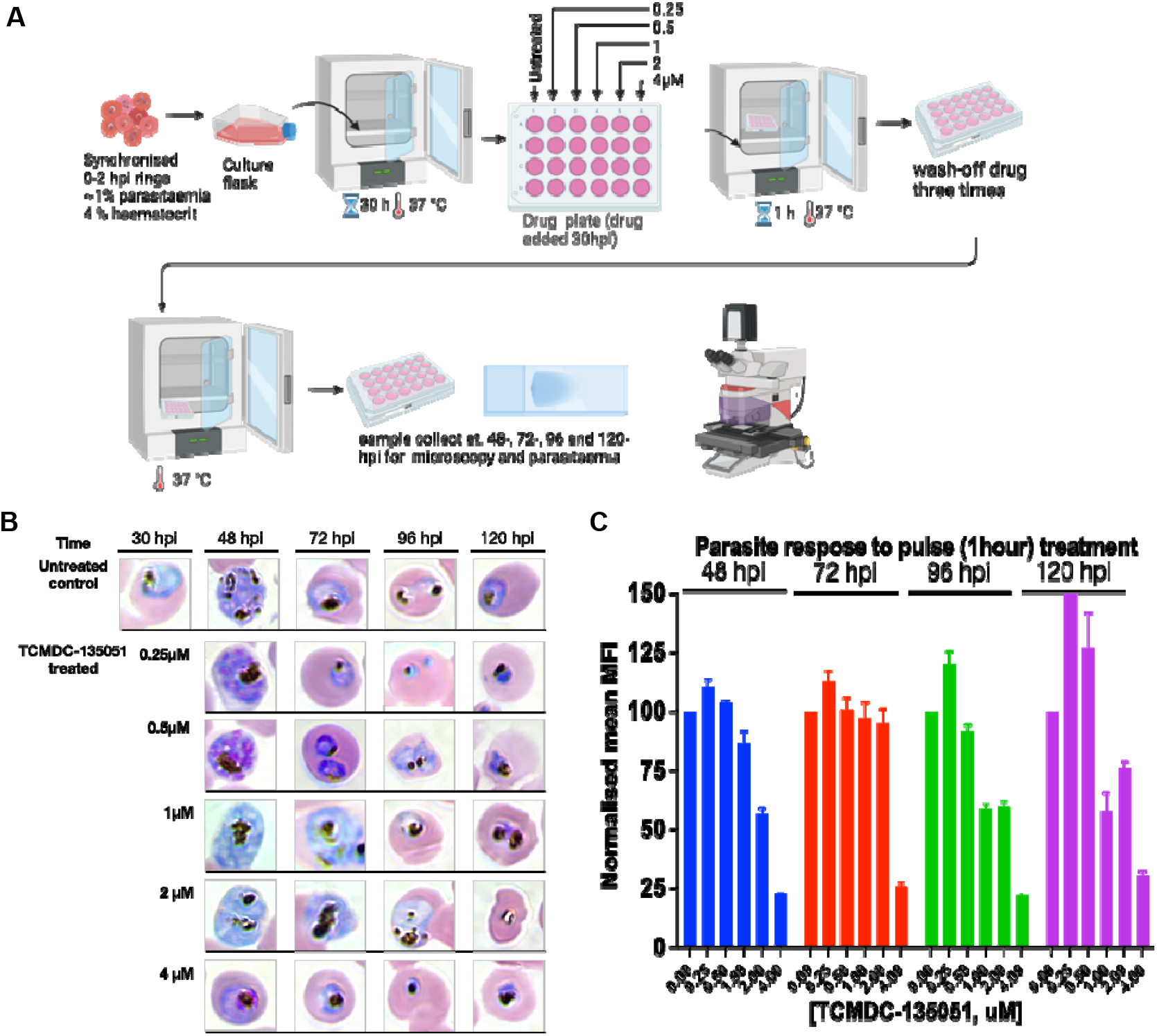
Schematic representation of workflow for pulse treatment, and Giemsa images. **A)** Step-by-step workflow for growing synchronised parasites to the appropriate treatment time of about 30-hpi. The parasites were then exposed to various drug concentrations and incubated for 1-hour at 37°C before been washed thrice to remove drug pressure. Samples were taken at 48-, 72-, 96-, and 120hpis. **B)** Images of Giemsa-stained thin smears collected at 48-, 72-, 96 and 120hpi. The images are representative of three independent experiments in triplicates **C)** Parasitaemia readout following incubation for up to 120-hpi. Error bars represent standard error of three independent experiments in triplicates (N=9).

In these experiments, parasites exposed to TCMDC-135051 concentrations between 0.25 - 1 μM for 60 minutes show no significant morphological impact on trophozoite progression through the parasite life cycle. The parasites matured to schizonts and produced invasive daughter merozoites. These parasites were maintained in culture, and no deleterious effect on development, maturation and re-invasion of fresh erythrocytes were detected **(Fig. 1B and C)**.

At higher concentrations (2 and 4μM), however, trophozoites-stage parasites showed significant morphological changes, and appeared visibly shrunken at 96-hpi **(Fig. 1B)**. This corresponds to the observed decreased parasitaemia **(Fig. 1C)**. Based on this evidence, exposure of trophozoite-stage parasites to 1 μM TCMDC-135051 inhibitor for 60 minutes was chosen as the suitable condition to investigate the effects of *Pf*CLK3 inhibition on gene transcription and RNA splicing. For parasite treatment, sampling and RNA preparation, see supplementary figure, Fig. S1.

### Inhibition of *Pf*CLK3 results in disruption of essential gene transcription

The inhibition of *Pf*CLK3 significantly impacted gene transcription, resulting in the downregulation of 218 transcripts and up-regulation of seven in wild type 3D7 parasites (WT) **(Fig. 2A, Table S1)**. In contrast, treatment of the TCMDC-135051 resistant mutant parasite strain, G449P, exhibited a markedly reduced transcriptional response to treatment **(Fig. 2B, C, Table S1)**. Importantly, under control conditions (i.e. without drug exposure) there were no detectable differences in gene transcription between the WT and G449P mutant parasites, indicating that the G449P mutation alone has no effect on transcription under no drug conditions **(Fig. S3**). Overall, these findings strongly support the conclusion that TCMDC-135051 specifically targets *Pf*CLK3 to modulate gene transcription.

**Figure 2.**
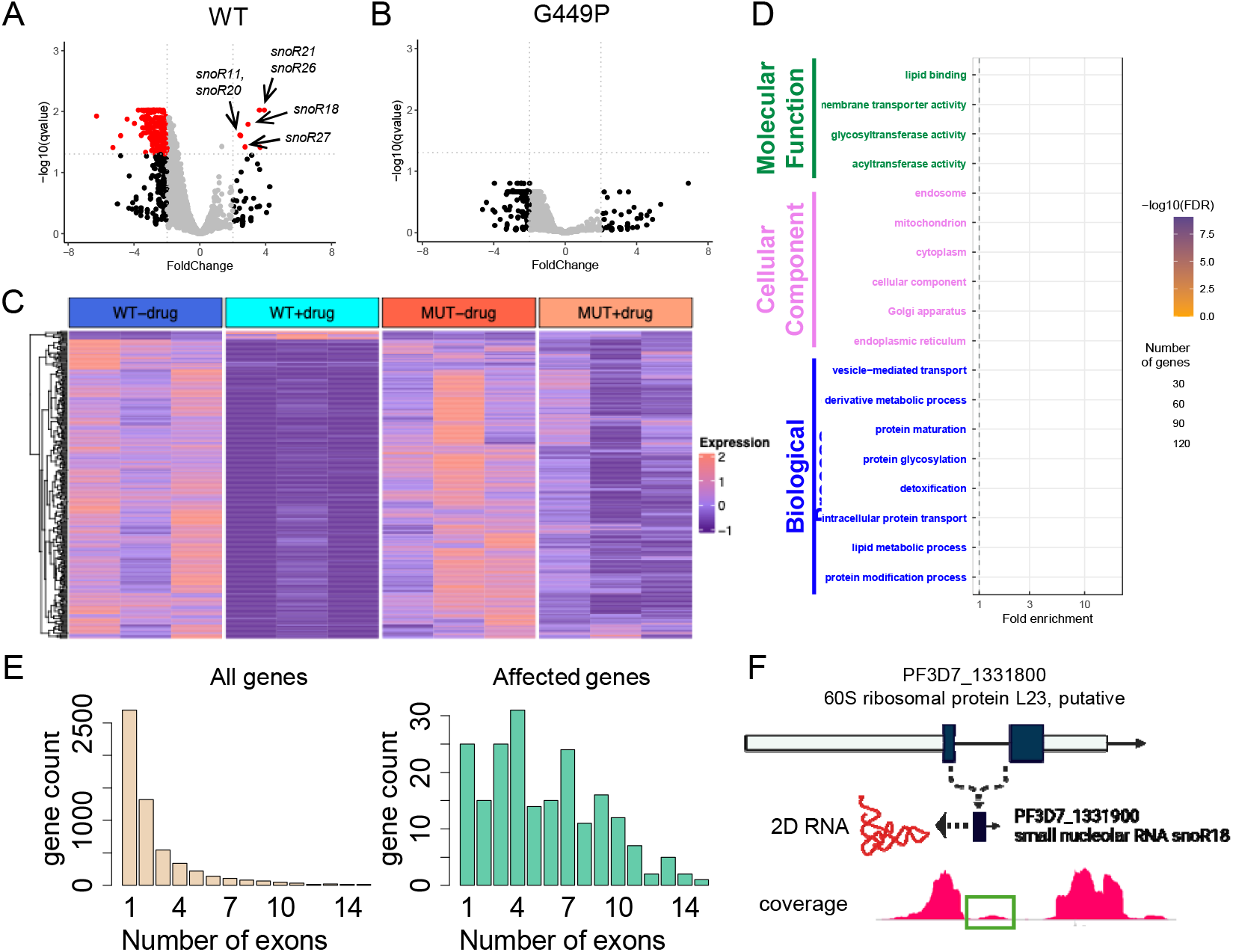
Gene expression changes in response to PFCLK3 inhibition. **(A-B)** Volcano plots showing differential gene expression in WT (A) and G449P (B) parasites under TCMDC-135051 treatment. Red=genes that reached the threshold of differential expression (log2 fold change > 2 and corrected p-value < 0.05); **(C)** Heatmap on of 214 genes differentially expressed in repones to *Pf*CLK3 inhibition in WT parasites showing their expression in both WT and G449P samples. Each row represents a gene. **(D)** GO term enrichment in differentially expressed genes (with multiple testing corrected P-value < 0.05). **(E)** Distribution of genes according to number of exons in whole transcriptome (left) and amongst differentially expressed genes (right), **(F)** the production of small nucleolar RNA (snoR18) from the intron of 60S ribosomal protein and its coverage in one of the samples.

Ontology analysis of down-regulated genes upon *Pf*CLK3 inhibition revealed enrichment in pathways critical to multiple essential parasite processes. This included genes involved in protein translation, intracellular trafficking and post-translational modification as well as genes regulating mitochondrial function and metabolic processes (**Fig. 2D**). The most striking feature, however, is that, of the down-regulated genes, 92.5% contain introns. Since approximately 50% of *P. falciparum* genes possess introns (10), this is a significant over-representation (P=2.2E-16) of intron containing genes. The genes containing multiple introns also appear to be more sensitive to *Pf*CLK3 inhibition (**Fig. 2E**). Finally, although most affected transcripts are down-regulated following treatment, seven transcripts are up-regulated in response to treatment. Notably, six of these encode for small nucleolar RNA (snoRNA) species associated with controlling RNA processing and modifications; and are derived by processing of the introns from larger genes (typically those encoding ribosomal proteins) (**Fig. 2F, S4**) suggesting the possibility of compensatory mechanisms acting to limit the impact of TCMDC-135051 on RNA metabolism. Together these data prompted a more direct examination of the impact of *Pf*CLK3 inhibition on RNA splicing.

### Effects of *Pf*CLK3 inhibition on RNA-splicing

A visual inspection of the coverage across the gene length revealed that, while clear intron boundaries were present in untreated parasites, splicing in treated WT samples was significantly affected, with multiple reads spanning introns. In untreated WT samples, approximately 90% of each transcript appeared to be correctly processed, whilst in treated WT samples, proper splicing dropped to as low as 10% for some junctions (**Fig. 3A**). In contrast, G449P parasites showed little to no defects in differential gene splicing between treated and untreated conditions for the same transcripts.

**Figure 3.**
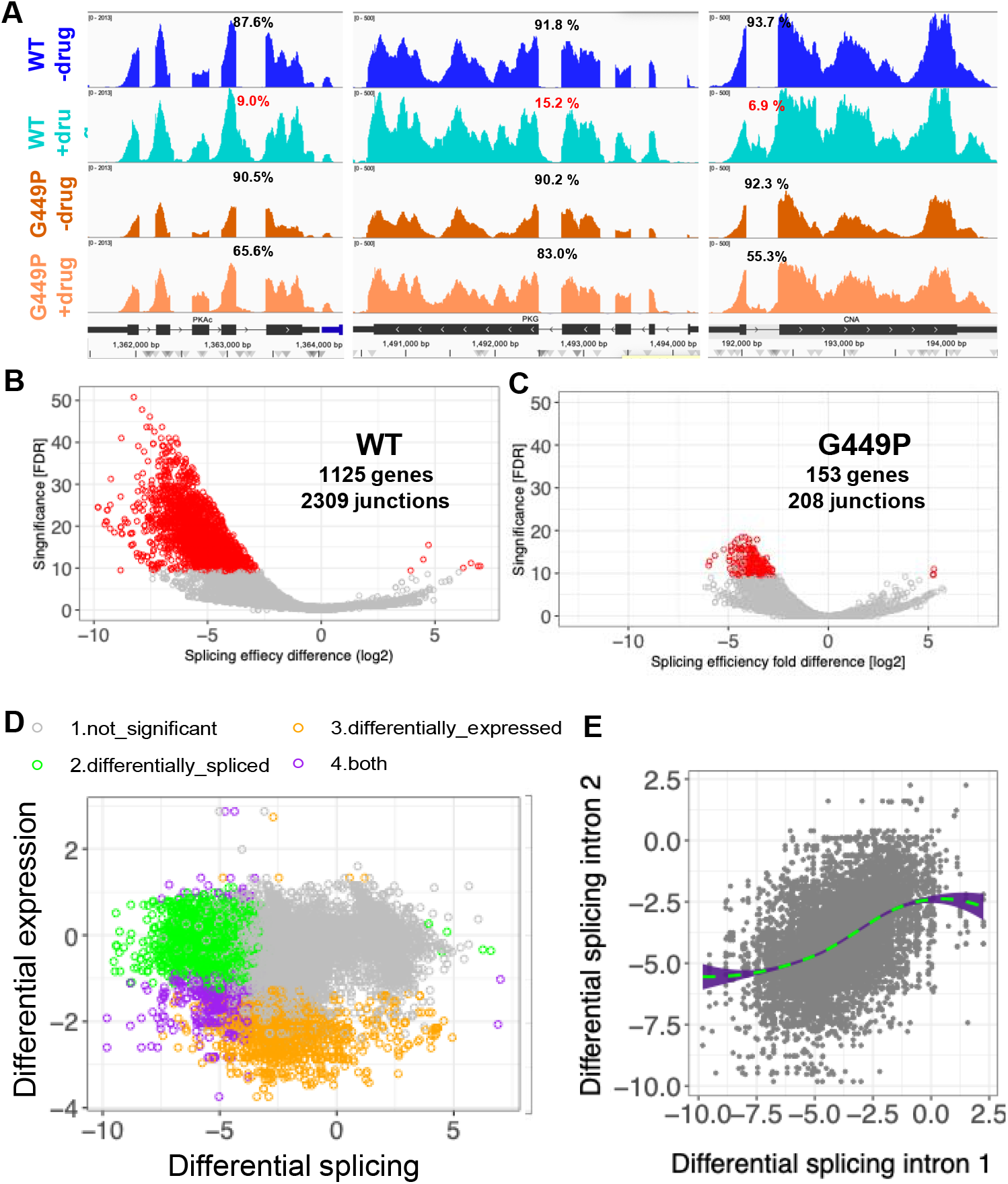
Analysis on global splicing landscape under the influence of TCMDC-135051 treatment. **(A)** Coverage plots showing disturbed intron processing in WT and G449P samples. As shown, splicing of introns in WT were efficiently completed in untreated WT in the range of about 90% compared to treated samples at between 7% and 15% for the same regions for PKAc, PKG and CNA genes, reducing splicing by more than 85%. This demonstrates the strong role of *Pf*CLK3 in splicing. However, in G449P parasites, inhibition reduces splicing only by a maximum of 37% for PKAc, PKG and CNA. For each condition only one of the 3 replicates is shown. **(B-C)** Volcano plots showing differential splicing in WT and G449P parasites upon TCMDC-135051 addition. Each point represents a spicing junction. Significantly differentially spliced genes (>25% in splicing difference and FDR value <0.001) are shown in red. **(D)** Comparison between differential splicing and differential expression. Fold difference in splicing efficiency in WT parasites after TCMDC-135051 pulse vs fold difference in the expression of the gene in which the junction is located. Points coloured according to the FDR values for both differential splicing and expression. **(E)** the effect of the drug on the junctions within the same transcript. Differential splicing between each intron pair located within the same transcript. Trend line shows a local polynomial regression model prediction of differential splicing of intron 2 based on intron 1

Global analysis of the splicing efficiency identified 2309 junctions across 1125 genes significantly affected by the action of the compound (**Fig. 3B-C**). Further investigation of the nature of this defect revealed that there is little correlation between expression level of the gene and splicing defect (**Fig. S5A**), ruling out potential artefacts of data analysis. While many genes affected on splicing level were also differentially expressed (n=163), the two phenomena were largely independent, with several genes correctly spliced but differentially expressed or the opposite (**Fig. 3D)**. As for differential expression, the G499P mutant was largely unaffected with only 208 junctions across 153 genes compared to WT, confirming the leading role of CLK3 in this phenomenon (**Fig. 3C)**.

Affected junctions were further analysed to assess the selectivity of this phenomenon, given that the majority of the 10,566 junctions remained unaffected. The correlation between the disturbance of junctions within the same transcripts suggests that the phenomenon is transcript specific rather than intron-specific (**Fig. 3E)**, although variation was still observed within the same transcript. All junctions were canonical major spliceosome junctions (of the GU/AG type) with no enrichment of any particular motifs in either the 200 bp flanking regions or the introns themselves (**Fig. 4A)**. The analysis of expression profiles of affected genes further reveal that distribution disproportionally affect genes with peak expression in trophozoite and schizont stages, consistently with the timing of the drug pulse delivery (**Fig. 4B)**. However multiple genes with stage-specific expression in other life stages were detected, suggesting that the drug may cause similar defects in merozoite or ring stages. Taken together, these findings suggest that the CLK3 efficacy is does not corelate with sequence specific binding or with expression timing but may instead depend on other factors such as RNA transcript turnover or translation speed.

**Figure 4.**
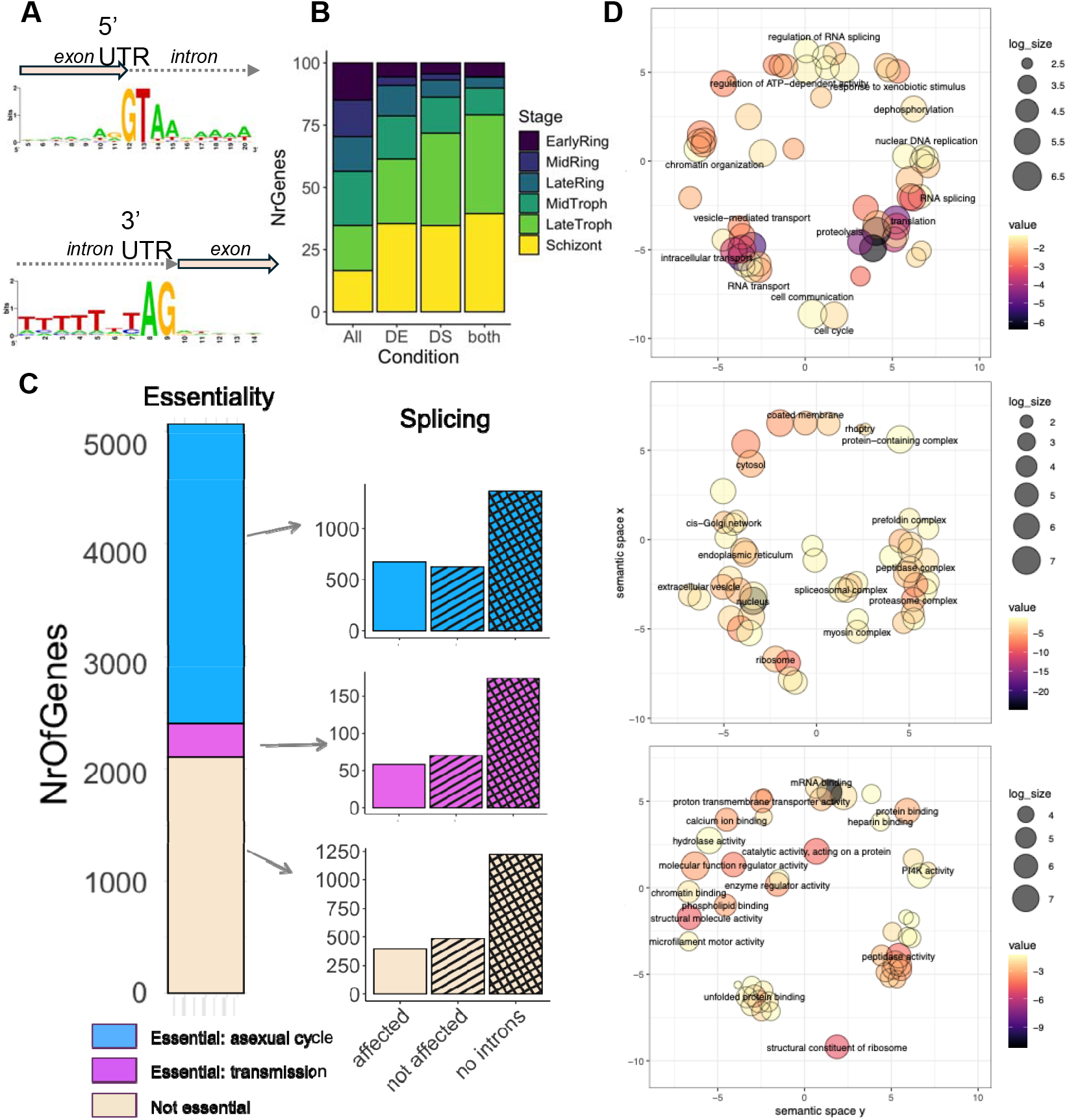
Analysis of junctions misplaced upon CLK3 inhibition in schizonts: **(A)** Affected junctions’ base composition at the 5’ and 3’ end, **(B)** timing of expression of differentially expressed and differentially spliced genes according to Painter et al, 2017. All=all transcripts in the genome, DE= differentially expressed, DS =differentially spliced, both = differentially spliced and differentially expressed; **(C)** Essentiality status of affected genes (using Zhang et al., 2019 for asexual replication and combination of Russell et al, 2023 and Stanway et al., 2019 for sexual development. **(D)** GO term analysis of the processes affected by aberrant splicing. Semantic space for Biological Process (BP), Cellular Components (CC) and Molecular Functions (MF) are shown with most informative terms printed. Full list of affected processes and genes contributing to them found in table S1.

The implications of splicing inhibition for the parasite biology are very significant. The functions of affected transcript spanned a wide range of pathways, with a total of 163 terms impacted (Fig. 4D, Table S3) - including essential processes for parasite survival such as host entry (GO:0044409), nuclear DNA replication (GO:0033260) and translation (GO:0006412). Among the most affected process are multiple aspects of RNA metabolism, including mRNA transcription (GO:0006367), splicing (GO:0008380) and transport (GO:0050658). This potentially reflects an increased turnover of the machinery in response to multiple incorrectly spliced transcripts, which paradoxically may lead to even greater dysregulation.

The analysis of the essentiality status of affected genes showed that at least 576 were essential for asexual replication, and 54 were essential for later stages such as gametocytogenesis or mosquito transmission (Fig. 4C). This suggests that splicing inhibition would lead to both a reduction of asexual parasitaemia and decreased transmission.

### PCR confirms inhibition of splicing in selected transcripts

To validate the splicing disruption observed from the bioinformatic analysis, we selected genes with introns that were dysregulated in the presence of the *Pf*CLK3 inhibitor. We then investigated whether these disrupted splicing events could be detected by PCR amplification in both WT and G449P parasites. Primers were designed to amplify both spliced and unprocessed transcripts (**Fig. 5A**), ensuring the detection of splicing defects involving a single intron (PF3D7_0321400), or multiple introns (PF3D7_0607600).

**Figure 5:**
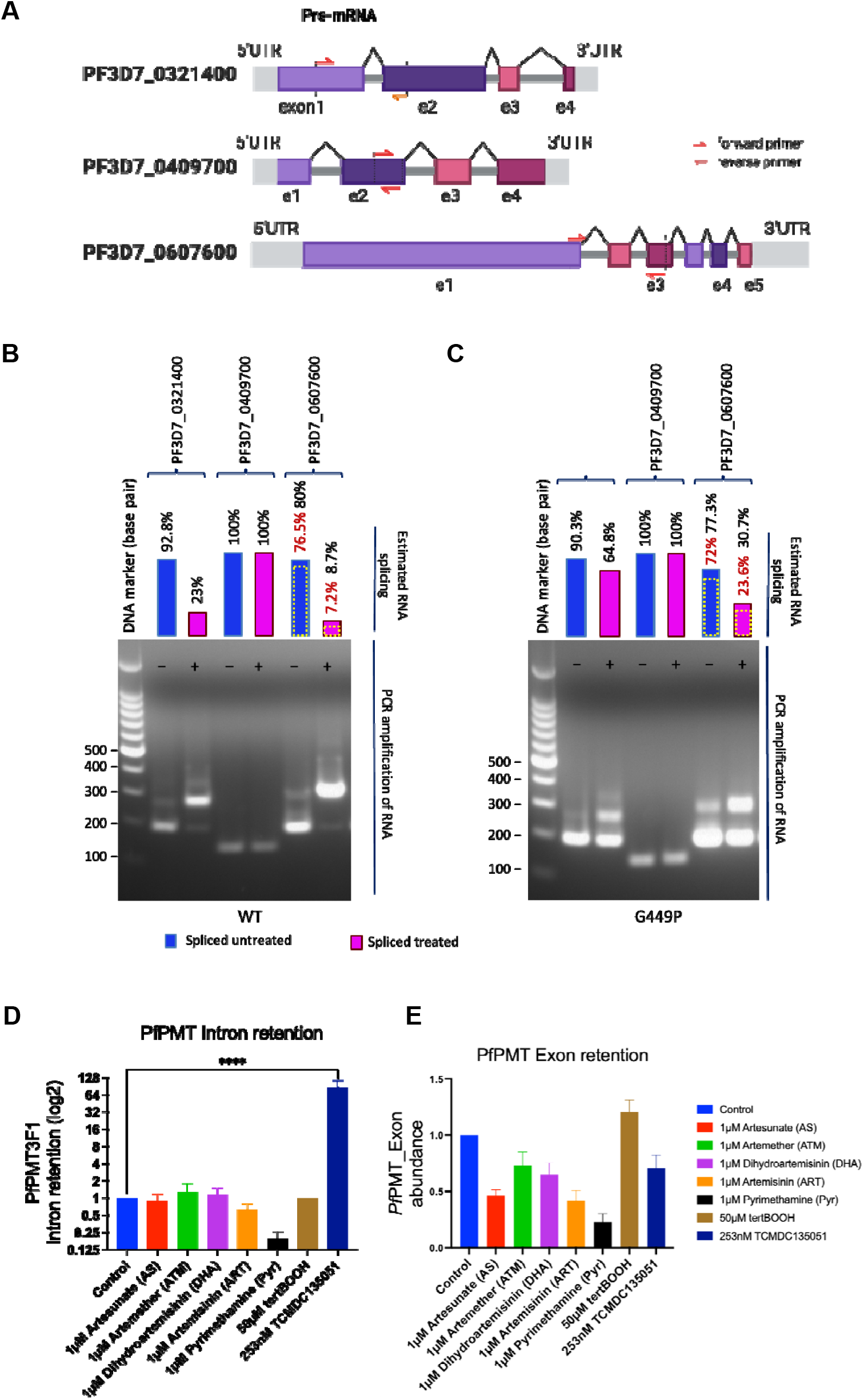
**A)** Schematic diagram of selected genes with disrupted splicing in WT parasites when treated with the *Pf*CLK3 inhibitor. Oligonucleotide PCR primers designed flanking selected predicted single introns that might be alternatively spliced as in PF3D7_0321400; an exonic region that would not be spliced, as in PD3D7_0409700; or spanning across three exons (flanking two introns) as shown with the gene PD3D7_0607600. **B and C)** PCR amplification of target regions showing splicing levels in the absence/presence of inhibitor in WT and G449P parasites. Top panel – shows splicing completing as detected from the RNA-seq data confirming alternative splicing in untreated (-, blue bar) and TCMDC135051 treated (+, pink bar) WT and G449P parasites. While splicing efficiency overlap between WT and G449P in untreated samples, there are obvious difference in splicing comparing treated WT and G449P parasites. **D)**. Comparing intron retention between TCMDC-135051 treated WT parasites to other commonly artemisinin and its derivates. Higher intron retention in TCMDC-135051 confirms splicing inhibition. (Error bars represent standard error of 3 independent experiments in triplicates. **E)**. Exon retention was evaluated but no significant difference was observed.

In the absence of the inhibitor, splicing efficiency was comparable between WT and G449P parasites (e.g. 76.5% and 80% respectively). However, in the presence of *Pf*CLK3 inhibitor, splicing efficiency in WT parasites was drastically reduced to 7.2% and 8.7% for first and second junctions respectively. In contrast, G449P parasites maintained high splicing efficiencies of 23.6 and 30.7% at the same junctions – representing a three-fold better splicing efficiency. These results further indicate that the differential splicing observed is directly attributable to the role *Pf*CLK3 in regulating splicing (**Fig. 5B and C**).

Since splicing efficiency is not typically assessed in standard drug assays, we employ a qPCR assay to examine the behaviour of one of the affected spliced junctions under a range of commonly used sub-lethal antimalarial drug treatments. These included the current first-line therapy components — artemisinin and its derivatives used in combination therapy (artesunate, artemether, and dihydroartemisinin)—as well as pyrimethamine, tertBOOT and TCMDC-135051. Intron retention was quantified following parasite exposure to each drug **(Fig. 5D)**. As expected, parasites treated with the *Pf*CLK3 selective inhibitor TCMDC-135051 exhibited significantly higher intron retention, consistent with splicing disruption. In contrast, exon expression remained unchanged across all treatments (**Fig. 5E**).

## Discussion

*Pf*CLK3 has been previously identified as a promising drug target, with its inhibition offering the potential for curative, transmission blocking and prophylactic effects (1). However, the molecular mechanism of action of *Pf*CLK3 inhibitors and their impact on parasite biology remained unclear. In this study, we demonstrate that *Pf*CLK3 inhibition has a broad and specific impact on parasite transcriptome, leading to a substantial reduction in intron processing. This disruption affects specific introns and splice junctions, ultimately resulting to parasite death. The *Plasmodium* genome is highly compact, characterised by a dense arrangement of open reading frames (ORFs) and frequently overlapping untranslated regions (UTRs) of neighbouring genes. Despite this compactness, approximately half of the genes in *Plasmodium falciparum* (52.5%) contain introns - often multiple – making the parasite highly reliant on an efficient splicing machinery to sustain essential cellular functions. Additionally, several *Plasmodium* genes have been shown to produce alternative splicing isoforms (11). While the functional significance of most of these isoforms remained uncharacterised, it is plausible that the parasite utilizes alternative splicing to diversify its proteome, enabling fine-tuning of gene expression necessary for specific cell types across different developmental stages and host species. Inhibition of *Pf*CLK3 provides new insights into its regulatory role and importance in splicing in *P. falciparum* parasites.

The study presented here demonstrate that *Pf*CLK3-inhibition leads to widespread splicing dysregulation, affecting approximately 20% of the transcriptome and 22% of all junctions. This disruption results in large accumulation of unprocessed transcripts and proteins, ultimately causing parasite death. Interestingly, a short drug pulse - even at high concentration - had minimal immediate impact on parasite progression through the life cycle. Most parasites were able to successfully complete the generation of progeny merozoites and subsequent invasion. This suggest that parasites may persist for a period using pre-accumulated proteins or transcripts, or, by producing excessive amount of RNA allowing them to survive even when only a small fraction is correctly processed. It also implies that, in contrast to higher eukaryotic cells, *Plasmodium* may tolerate high level (>90%) of aberrant (nonsense) transcripts without immediate damage or functional consequences. The latter would be consistent with the fact that not many of these transcripts are actively degraded (as seen through differential expression analysis) and the fact that nonsense mediated transcript decay is not essential for *Plasmodium (12)*. These findings should be considered during further optimisation of *Pf*CLK3, including decisions regarding dosing regimens, partner drugs and treatment duration. Moreover, *Pf*CLK3 inhibitors could serve as valuable tools for studying mRNA and protein turnover dynamics across different parasite life cycle stages – an area that remains poorly understood.

Detailed analysis of the affected junctions revealed that despite many genes affected by CLK3 inhibition, the effect is not global since a lot of transcripts are processed correctly while others remained completely unprocessed. The fact that affected transcripts are expressed mainly in schizonts could suggest that the effect could be influenced by RNA turnover (i.e. only transcripts highly expressed and degraded within 1-hour window of the drug action are affected). However, even within the same transcript, processing efficiency varies according to different junctions (Fig. 3A, E), suggesting that some splicing events are more sensitive to inhibition than others. This is consistent with previous work performed on *Pf*CLK3 homolog in yeast (Prp4), which when inhibited, resulted in decreased splicing of only a section of introns, characterised by “weak” 5_′_ splice sites or suboptimal branch point sequences (13). Similarly, in *Toxoplasma gondii*, the inhibition of CLK3 homologue, *Tg*PRP4, resulted in disturbed splicing of second intron of many genes (14), also suggesting the selectivity of the function. The mechanistic basis of CLK3 selectivity at these junctions requires further investigation across different parasite life stages and under varying drug-pulse intervals. Identifying particularly sensitive junctions would deepen our understanding of *Plasmodium* biology and could enable the development of new assays, facilitating medium-throughput screening for additional splicing inhibitors.

In addition to advancing our understanding of *Plasmodium* biology, this work further validates *Pf*CLK3 as potential drug target for new antimalarials. To address the growing resistance to current front-line malaria treatments, next generation anti-malarials must act on novel targets and possess unique mechanism of action. Ideally, these compounds should be effective across multiple stages of the parasite life cycle - including the liver stage, offering prophylactic; the asexual blood stages, for curative action; and the sexual (gametocyte) stages to block transmission (15,16). The molecular mechanism of action of TCMDC-135051 supports the potential of *Pf*CLK3 inhibitors to meet these criteria.

*Pf*CLK3 inhibitors are currently being developed to meet the target product profile (TCP) defined by Medicines for Malaria Venture (MMV), specifically for a cure (TCP-1) and chemoprevention (TCP-2) (15). Although developing highly selective protein kinase inhibitors is inherently challenging, availability of a high-resolution X-ray crystal structure of *Pf*CLK3 in complex with TCMDC-135051 has enabled a structure-based drug design approach. This strategy is yielding highly selective next generation inhibitors. Combined with their multi-stage and cross-species activity, *Pf*CLK3 inhibitors satisfy key requirements for a next-generation anti-malarial therapy with the potential to deliver a radical cure.

## Methods

### Parasite culture and drug assays

Asexual blood stage *P. falciparum* 3D7 parasites were cultured at 37 ° C in RPMI (Roswell Park Memorial Institute) media supplemented with 0.5% Albumax II (ThermoFischer Scientific) as a serum substitute, 0.292g L-glutamine, 0.05g hypoxanthine, 2.3g sodium bicarbonate, 0.025g gentamicin, 5.957g HEPES and 4g dextrose in 1 litre RPMI. Parasites were grown at 4% haematocrit using blood from O+ male anonymous donors, provided through the UK Blood and Transfusion service. Parasite cultures were maintained in a parasite gas atmosphere (90% Nitrogen, 5% Carbon dioxide, 5% oxygen) (16). Parasitaemia is routinely monitored by Giemsa staining of methanol-fixed air-dried thin blood smears visualised by light microscopy. Parasite cultures were synchronised by isolating mature schizont-stage parasites on a cushion of 60% Percoll (GE Healthcare). Purified schizonts were incubated in complete media at 37°C with fresh RBCs for 1-4 hours in a shaking incubator to allow new merozoites to invade fresh erythrocytes. Remaining schizonts were removed by a second Percoll purification leaving tightly synchronised ring-stage parasites.

### Establishment of drug concentrations, RNA Sample collection and preparation

To determine the optimal sublethal concentration of TCMDC-135051 that would allow assessment of its effect on parasite pre-mRNA processing and splicing without killing the parasites, doubly synchronized ring-stage parasites (0–3 hours post-invasion, hpi)— WT and G449P—at 1% parasitaemia and 4% haematocrit (HCT) were aliquoted into a 96-well flat-bottom plate and cultured up to the late trophozoite stage (30 hpi).The parasites were then exposed to varying concentrations of TCMDC-135051 (0, 0.25, 0.5, 1, 2 and 4μM) for 1-hour at 37 °C. Drug pressure was removed by washing the cultures three times with complete media and parasites incubated further up to 120 hpi. Giemsa-stained thin smears were collected at 30- and 48hpi, and every 24 hours up to 120-hpi to monitor parasite replication, (Fig. 1A-C). As proxy for parasitaemia, Sybr green assay was used to measure DNA staining as direct measure for parasitaemia. Assessment of parasite morphology and parasitaemia indicates that up to 1μM drug exposure had little effect on parasite replication and was used in subsequent steps.

To collect samples for RNA-sequencing, doubly synchronised WT and G449P parasites were cultured for 30 hpi (-/+2 hrs) and then treated with 1μM TCMDC-135051 inhibitor or vehicle, (DMSO) for 1 hour at 37°C. Parasitised erythrocytes were harvested by centrifugation and stored in TRIzol in −80°C until RNA extraction. Parasite total RNA was extracted using the Qiagen RNeasy Mini Kit according to the manufacturer’s instructions. RNA quantity and quality were measured by NonoDrop 2000 (Thermo Fisher Scientific). Both treatment and controls were run in triplicates under the same culture conditions. Complementary DNA, cDNA was generated using QuantiTect Reverse Transcription Kit according to manufacturer’s instruction.

### RNA-seq data generation

Purified RNA quality was checked using Agilent Bioanalyser and Nanodrop spectrophotometer. Illumina TruSEQ RNA library prep and sequencing reagents were used following the manufacturer’s recommendations using polyadenylate-selected transcript selection. The samples were paired-end multiplex sequenced (2x 150 base pairs) on the illumina HiSeq 2500 platform, and at least 30 million reads were generated for each sample. The quality of the raw sequencing reads was assessed using FastQC software.

### Data analysis

#### Pre-processing

The RNA-seq reads (FASTQ) was mapped against Plasmodium falciparum reference genome (downloaded from PlasmoDB v. 56) using splice-aware aligner hisat2 (v. 2.2.1) (17). Resulting BAM files were sorted, indexed and filtered for the mapping quality (only reads with MAPQ>10 were kept) using Samtools suite (v. 1.17) (18) and visually inspected using IGV genome browser (19).

### Differential gene expression analysis

The number of reads mapping against each of the annotated genes was calculated with *featureCounts* function from subread software package (v.2.0.0) using the genome feature file (gff) corresponding to the used genome version. R software with edgeR function package was used to normalise the data, estimate dispersions and fit generalised linear model to the dataset with default parameters, as specified in the software vignette. Pairwise contrasts were set up between the two lines in the same conditions (G449P vs WT) or between one line in two different conditions using r quasi-likelihood (QLF)-test. A log2-fold change cut off in gene expression of >2 and P-values of statistical test with Bonferoni multiple testing correction of <0.05 were used to call differentially expressed genes.

### Differential splicing analysis

Conventional tools used for splicing analysis based on isoform usage quantification proved suboptimal for the *Plasmodium* genome, likely due to its high gene density and uneven exon coverage. To address this, we developed a custom approach method to quantify differences in intron processing across parasite lines and treatment conditions (Fig. S1). Within each gene we considered all the candidate splice junctions, i.e. all the junctions in the reference annotation from PlasmoDB, plus all the genomic intervals flanked by the canonical splice sites GT/AG. For each splice junctions in each RNA-Seq library, we quantified two types of reads: those supporting splicing - i.e. reads split by Hisat2 aligner (17) to bridge the two adjacent exons - and those supporting intron retention - i.e. reads partially overlapping an exon and the adjacent intron. This quantification was performed using featureCounts (20) and custom scripts (see below). Differential splicing is the difference in the proportion of spliced vs unspliced reads in one condition versus another condition. To assess this difference, we applied the method developed for differential methylation (21), since effectively methylated vs unmethylated counts at a cytosine are equivalent to spliced vs unspliced counts at a splice junction. In brief, the method applies the generalised linear model (GLM) strategy implemented in edgeR package (21) where the vector counts (spliced and unspliced) at a junction is modelled as a GLM with negative bionomial distribution with covariates being the treatment effect and the count type (spliced or unspliced). Differential splicing is then the interaction effect between treatment and count type. To reduce statistical and biological noise we tested only junctions accumulating at least 15 reads in all the libraries from at least one condition. This workflow is available as a Snakemake pipeline at https://github.com/glaParaBio/denovo-detection-of-differential-splicing.

For each pairwise comparison, a *DGEList* object was created comprising the counts from relevant libraries, along with sample information and library sizes, which were calculated by summing the splice and unspliced reads counts for each sample. The design matrix was generated according to [ref.] and used for dispersion estimation. A negative binomial generalized log-linear model was fitted to the read counts of each intron, and modified likelihood ratio test (LRT) were conducted using either the strain or drug treatment as a contrast condition. The resulting table containing log-fold changes of the proportion of reads spliced, and P-values with Bonferroni correction were generated and used to filter the differentially spliced introns. Introns with >25% changes in splicing proportions between the conditions and q values <0.001 were considered as differentially spliced/retained. Visual inspection of selected junctions was performed by generating *.bw files using deeptools v. 3.3.2 and viewing them with Integrative Genomics Viewer v 2.3.91.

### GO terms analysis and comparison with existing datasets

GO terms analysis was carried out using topGO package in R. Terms containing at least two genes and P value of enrichment test <0.05 were considered significant. The number of exons in each gene and its analysis was calculated based on gff file and plotted using custom R script.

In order to compare to study the connection between the transcription time of the genes and effect of the compound on their splicing the dataset from *de novo* (ref) and steady state transcription were downloaded from repository and Chi-sequare test was used to test for the enrichment for the gene with peak of de novo transcription at the given stage (de novo data) or time point (steady state transcription).

### PCR amplification of cDNA

To show splicing disruption using standard PCR, selected of genes with introns were chosen. PCR primers were designed on exon regions flanking intron sites, (Fig. 5, Table S4). Amplifications of parasite cDNA were done using standard methods and the PCR product ran on 1% agarose gel to visualise amplified products.

## Data availability

The pipeline used for the differential splicing analysis is available at: https://github.com/glaParaBio/denovo-detection-of-differential-splicing. Illumina sequencing data have been deposited in the NCBI Gene Expression Omnibus (GEO) under the accession number GSE278703.

## Acknowledgements

The authors would like to acknowledge funding from the Gates Foundation (INV-039928 and INV-088781), Wellcome Trust and Royal Society (202600/Z/16/Z) and Singapore Ministry of Education grant (# MOE2019-T3-1-007)

---

**Figure S1.**
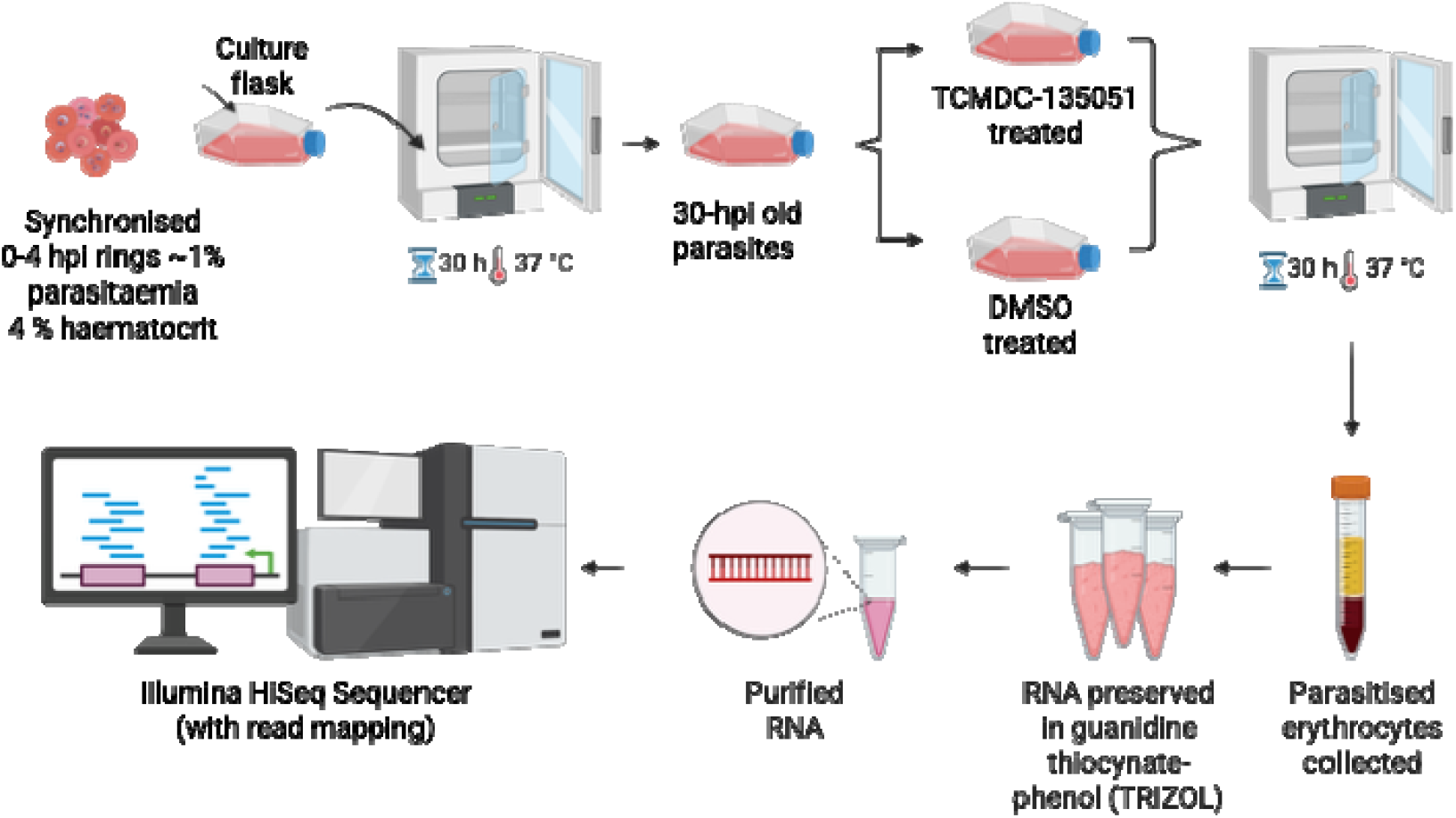
Schematic workflow for enriching parasites, RNA purification, and sequencing and data analysis.

**Figure S2.**
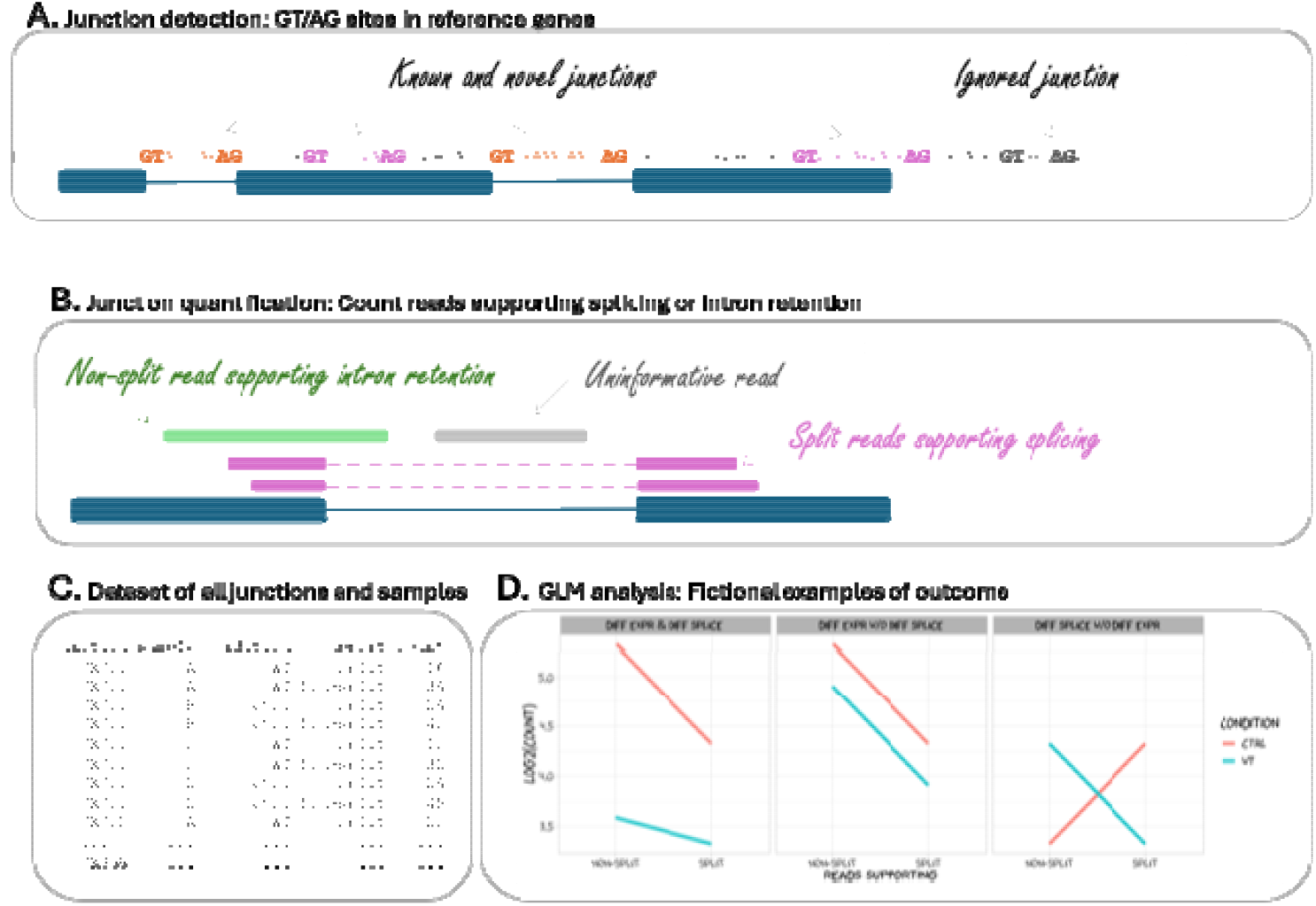
Strategy for identifying differentially spliced junctions in Plasmodium: **A)** Testable splice junctions are genomic intervals flanked by the canonical GT/AG sites; these junctions may or may not correspond to reference introns, but they must overlap reference genes. **B)** RNAseq reads split across junctions (purple) support splicing while reads straddled across junctions support intron retention (green); reads fully contained in introns, exons or intergenic regions are ignored. **C)** For each junction and for each library, compile a table of counts supporting splicing and intron retention. **D)** Differential splicing occurs when the difference between split and non-split counts is different between conditions (e.g. left and right plot, the plot on the left also has evidence of differential expression); the plot in the middle has evidence of differential expression but not of differential splicing.

**Figure S3.**
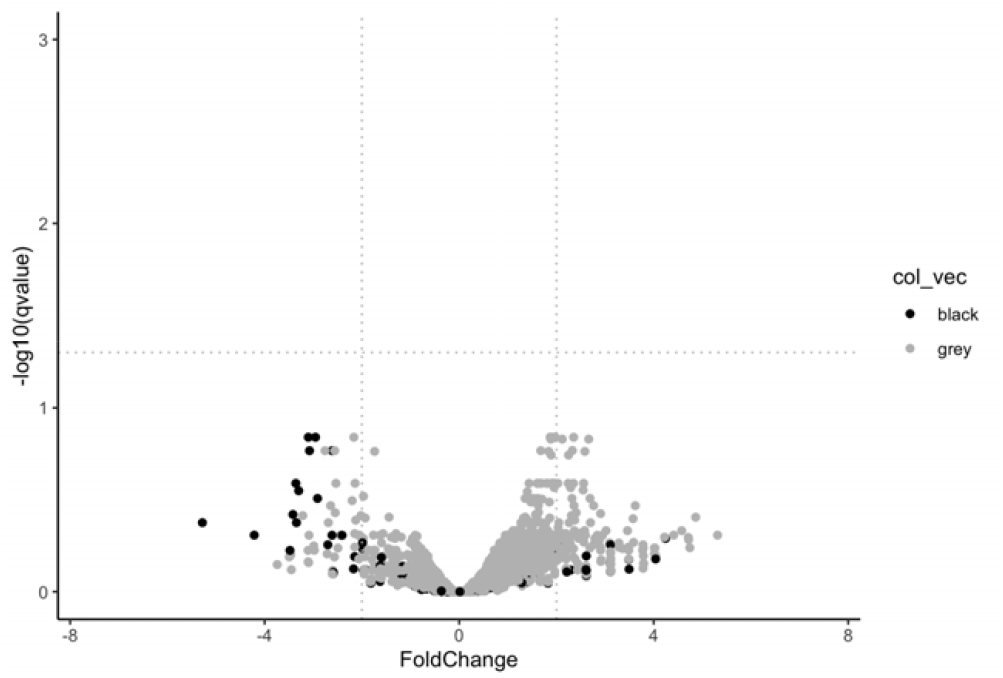
Differential expression between WT and G449P mutant.

**Figure S4.**
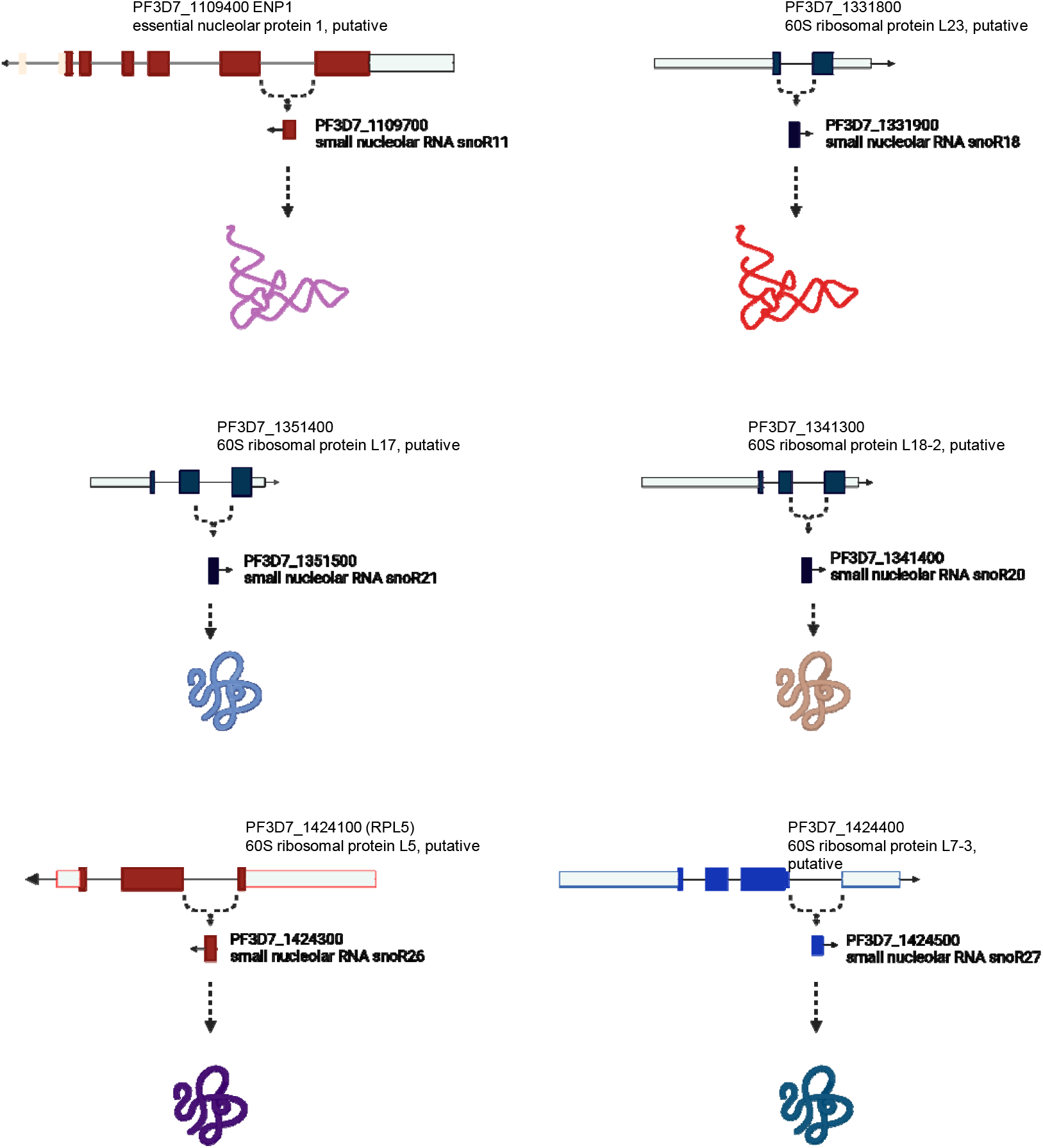
Schematic representation of snoRNAs affected by TCMDC-135051 treatment: (A-F) position of 6 out of 7 overexpressed snoRNA within the “host” genes.

**Figure S5.**
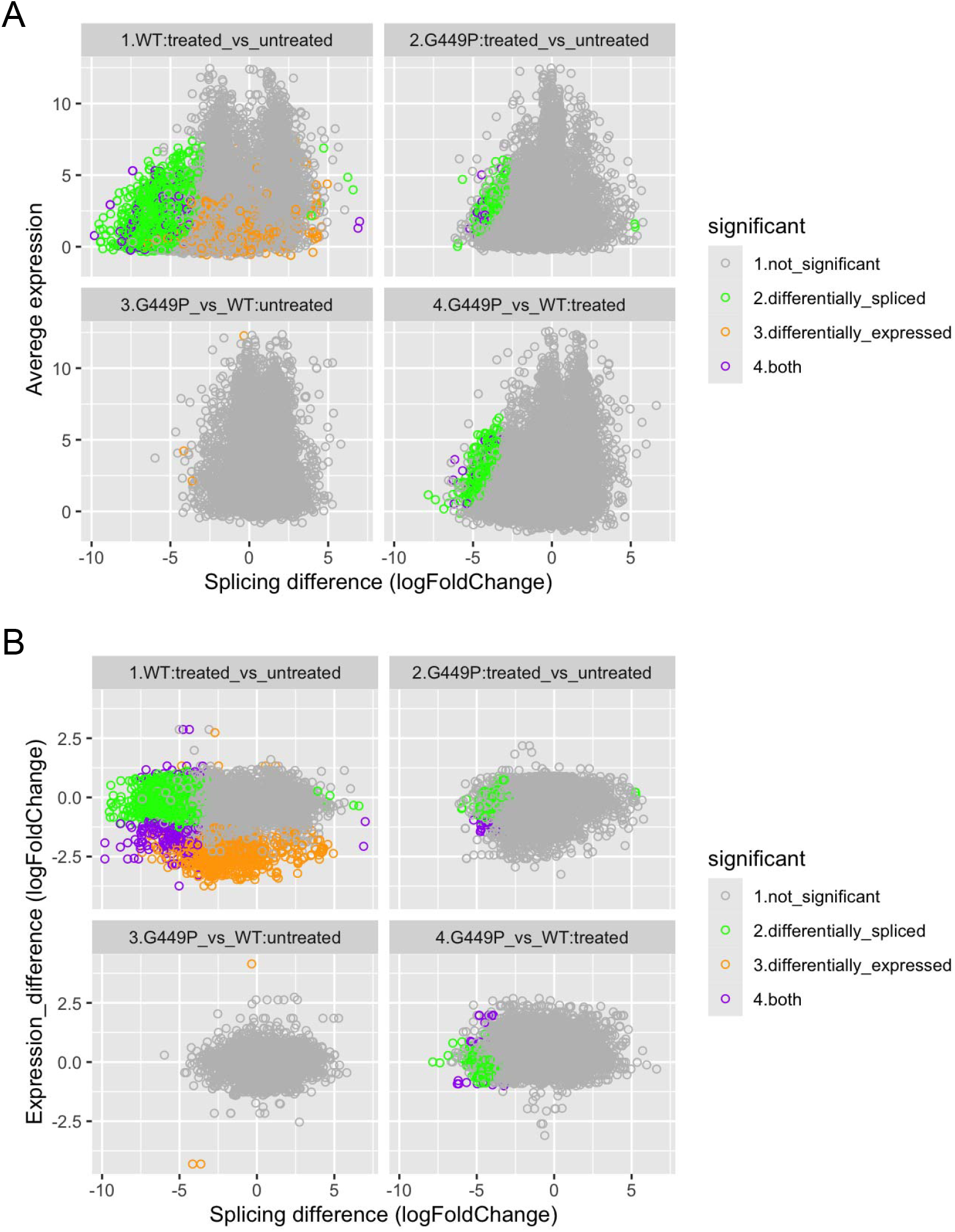
Differential splicing analysis across different conditions. **(A)**. Splicing efficiency difference across different expression levels **(B)** comparison of differential splicing and differential expression across different lines and conditions.

